# Refinement of efficient encodings of movement in the dorsolateral striatum throughout learning

**DOI:** 10.1101/2024.06.06.596654

**Authors:** Omar Jáidar, Eddy Albarran, Eli Nathan Albarran, Yu-Wei Wu, Jun B. Ding

## Abstract

The striatum is required for normal action selection, movement, and sensorimotor learning. Although action-specific striatal ensembles have been well documented, it is not well understood how these ensembles are formed and how their dynamics may evolve throughout motor learning. Here we used longitudinal 2-photon Ca^2+^ imaging of dorsal striatal neurons in head-fixed mice as they learned to self-generate locomotion. We observed a significant activation of both direct- and indirect-pathway spiny projection neurons (dSPNs and iSPNs, respectively) during early locomotion bouts and sessions that gradually decreased over time. For dSPNs, onset- and offset-ensembles were gradually refined from active motion-nonspecific cells. iSPN ensembles emerged from neurons initially active during opponent actions before becoming onset- or offset-specific. Our results show that as striatal ensembles are progressively refined, the number of active nonspecific striatal neurons decrease and the overall efficiency of the striatum information encoding for learned actions increases.

## INTRODUCTION

The nervous system possesses the remarkable ability to control animal behavior and to adapt this control to accommodate future actions and goals. This precise process involves the recruitment of diverse neuronal populations and circuits distributed throughout the nervous system. The striatum, the input nucleus of the basal ganglia, plays a critical role in motor control and is predominantly comprised of D1R-expressing direct pathway spiny projection neurons (dSPNs) and D2R-expressing indirect pathways SPNs (iSPNs) (Graybiel, 2005; Jin and Costa, 2010; Calabresi et al., 2014; Santos et al., 2015). The striatum receives convergent inputs from various sensorimotor areas, including the cortex and thalamus. Lesioning the motor cortex in animals trained to perform motor skills affects the behavior when lesions occur pre-training but not after they are well-trained (Lawrence and Kuypers, 1968; Kawai et al., 2015; Hwang et al., 2019; Dhawale et al., 2021). In contrast, lesioning the striatum not only prevents animals from learning new motor skills (Turner and Desmurget, 2010; Yin, 2010; Koralek et al., 2012), but also causes a variety of motor symptoms, including the disruption of the performance of previously learned motor skills (Lemke et al., 2019; Dhawale et al., 2021), delayed movement kinematics, altered coordination, postural abnormalities, and difficulty initiating movements (McHaffie et al., 2005; Markowitz et al., 2018).

Consequently, a central question in the neurobiology of action is how corticostriatal neuronal activity encodes actions and how these encodings change over time. Indeed, as an animal’s behavioral repertoire expands, it is imperative that precise neuronal plasticity mechanisms are engaged that facilitate integrating novel action encodings into ongoing neuronal population activity, minimizing encoding inefficiencies and potential conflicts between different action representations. Recent *in vivo* calcium imaging studies using longitudinal imaging of neuronal activity revealed that the activity patterns of individual neurons adapt over the course of motor learning (Komiyama et al., 2010; Huber et al., 2012; Masamizu et al., 2014; Peters et al., 2014; Peters and Komiyama, 2017). Head-fixed mice learning a lever push task leads to the formation of precise spatiotemporal activity patterns within the primary motor cortex (M1), with layer 2/3 neuron activity becoming more aligned to movement onsets and stable temporal sequences of layer 2/3 neuron activity emerging (Peters and Komiyama, 2017). In the striatum, motor learning results in neuronal activity becoming correlated with the initiation and termination of action sequences (Jin and Costa, 2010), and like the upstream motor cortex, striatal SPNs in the dorsolateral striatum (DLS) develop reproducible firing sequences that are specific to learned movements (Sheng et al., 2019; Maltese et al., 2021). Thus, within both M1 and DLS, neuronal ensembles emerge during motor learning whose activations are selective to specific features of movement.

These prior studies raise an important question – how are neuronal ensembles formed during learning in the striatum? In particular, the specific roles played by direct and indirect pathway ensembles across days of training, are not well-understood.

To address this question, we performed *in vivo* longitudinal 2-photon imaging of striatal Ca^2+^ dynamics as head-fixed mice learned to execute voluntary bouts of locomotion on a running wheel. By comparing periods where animals initiated bouts of locomotion (onsets) to periods where they terminated locomotion (offsets), we were able to isolate two distinct action states and their corresponding striatal activity, permitting us to investigate how striatal activity encodes (i.e., predicts) these distinct actions throughout the learning process. Furthermore, as animals progressively learned to self-locomote, we were able to study these striatal dynamics across days of learning. We found that both dSPNs and iSPNs are highly recruited during motion, but that their degree of motion-activation decreases throughout subsequent days of training. In particular, motion-nonspecific SPNs are progressively disengaged throughout learning. Concurrently, we observed that SPNs are progressively refined during learning such that by the late stage of training, the action-specific (onsets vs. offsets) ensemble sizes are smaller. Fate and regression analyses suggest that action-specific neurons do not emerge throughout learning but are instead present at the beginning of training. Furthermore, although the number of striatal neurons active during bouts of motion decreases throughout learning, the information content is preserved. In combination, these results suggest that learning coincides with disengagement of nonspecific SPNs, ultimately yielding refined, action-specific ensembles that more efficiently encode the new actions.

## RESULTS

### Longitudinal imaging of striatal Ca^2+^ dynamics during self-generated locomotion

The choice of behavioral paradigm dramatically impacts the identity and dynamics of the underlying neuronal activity being studied (Barnes et al., 2005; Markowitz and Datta, 2020). This is particularly true for the striatum, where lesions affect not only sensorimotor function, but also learning and memory of various goal-directed behaviors (Yin et al., 2004; Bailey and Mair, 2006; Liljeholm and O’Doherty, 2012; Rueda-Orozco and Robbe, 2015). With the aim of isolating the role that the DLS plays in encoding specific motor actions, instead of training mice on a motor task with complex reward and temporal contingencies, we placed head-fixed mice on freely moving running wheels, allowing them to execute bouts of self-generated locomotion. To longitudinally image the same SPN populations across days, an AAV encoding Cre-dependent GCaMP6s was injected unilaterally into the DLS of Drd1a-Cre or A2a-Cre mice to label dSPNs and iSPNs, respectively (Figure 1A,B). Mice have a natural tendency to locomote (Grillner, 2006; Arber and Costa, 2022), and our behavioral paradigm was designed so that the animals could freely learn to locomote in the novel head-fixed running wheel context. Indeed, across imaging sessions, mice progressively learned to locomote, as measured by the wheel’s average per-session velocity across days, as well as the proportion of session time spent running across days (Figure 1C,D). Importantly, our behavioral paradigm does not rely on rewards or task structure, as mice were allowed to freely execute self-generated bouts of locomotion without sensory cues or feedback, permitting us to directly investigate the relationship between striatal activity and the motor actions (i.e., motion onsets and offsets) being executed. With this approach, we were able to simultaneously measure bouts of locomotion and the Ca^2+^ dynamics of ∼160 striatal neurons in each animal across days as they learned to locomote freely on the wheel (Figures S1A and S2A,B).

**Figure 1.**
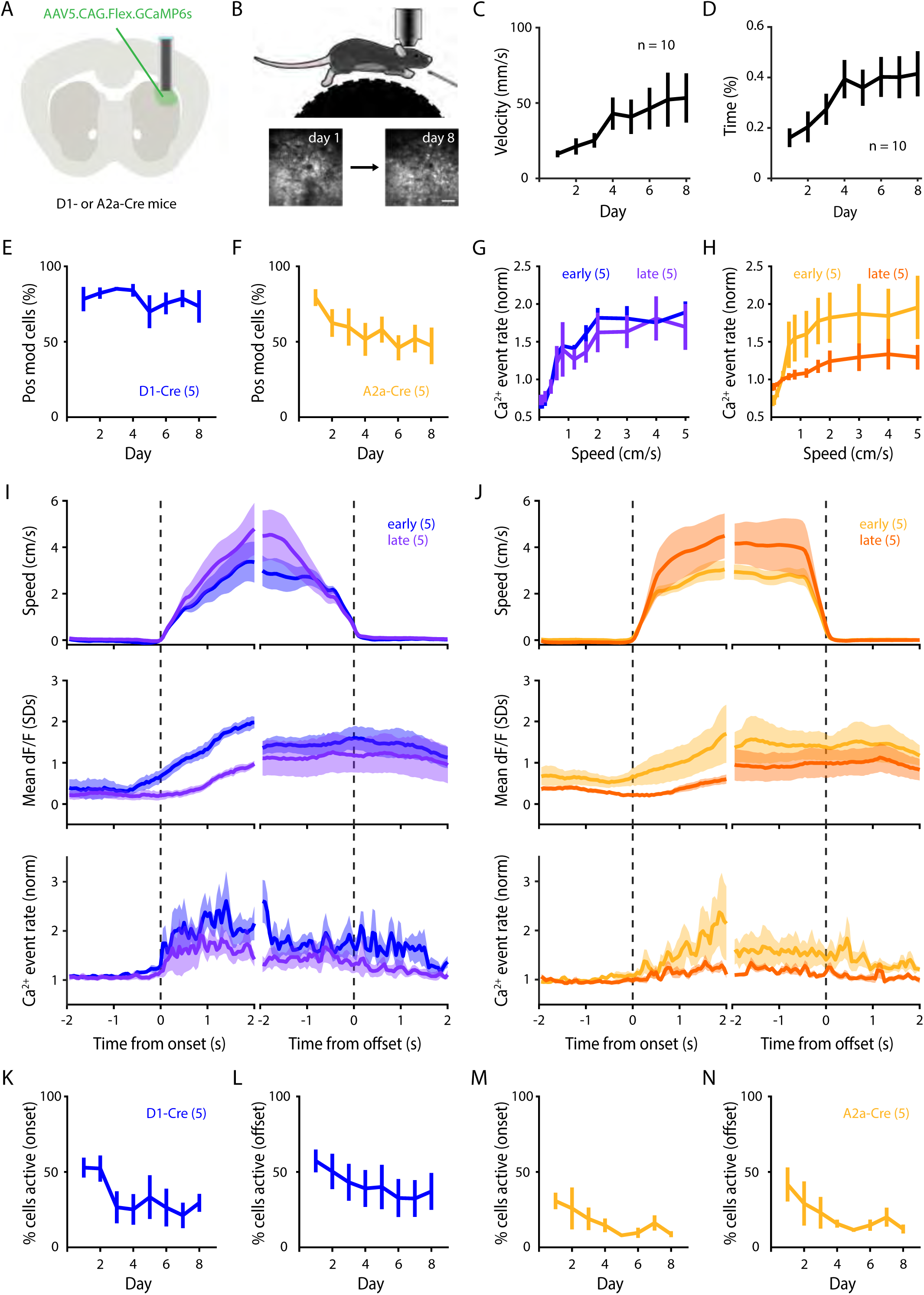
Striatal activation decreases throughout learning. **A,** Illustration of stereotaxic injection of AAV delivery of GCaMP6s into the DLS of D1-Cre or A2a-Cre mice followed by GRIN lens implantation. **B,** Schematic of 2-photon imaging during head-restrained, self-generated locomotion paradigm. Bottom: DLS longitudinal imaging, scale bar: 100µm. **C,** Average velocity of mice throughout each session (n = 10 mice; p = 0.030). **D,** Average time spent running (n = 10 mice; p < 0.0001) depicting a progressive increase in self-generated locomotion across days. **E, F,** Percentage of imaged neurons that were positively modulated during periods of locomotion (relative to periods of non-locomotion) for dSPNs (**E**; n = 5 mice; p = 0.475) and iSPNs (**F**; n = 5 mice; p = 0.030). **G, H,** SPN activity (event rate) plotted as a function of locomotion speed, averaged across ‘early’ and ‘late’ sessions for dSPNs (**G**; n = 5 mice; p = 0.971) and iSPNs (**H**; n = 5 mice; p = 0.011). **I, J,** Left: dSPN summary, right: iSPN summary. Top: average animal locomotion speed centered around motion onsets and offsets. Middle: average standardized dF/F activity of imaged SPNs centered around motion onsets and offsets. Bottom: average Ca^2+^ event rate (normalized) of imaged SPNs centered around motion onsets and offsets. All analyses compare average between ‘early’ and ‘late’ sessions. **K-N,** Across-day percentages of dSPNs active during motion onsets (**K**, n = 5 mice / 8 days / 298 onsets; p = 0.003), offsets (**L**, n = 5 mice / 8 days / 314 offsets; p = 0.005), iSPNs active during motion onsets (**M**, n = 5 mice / 8 days / 320 onsets; p = 0.008), and offsets (**N**, n = 5 mice / 8 days / 320 offsets; p = 1.343e-04). Data are mean ± SEM. Statistical significance was assessed by repeated measures 1-way ANOVA with multiple comparisons (**C-F**), 2-way repeated measured ANOVA with multiple comparisons (**G, H**), linear mixed-effects modeling (**K-N**).

### Striatal activity progressively decreases throughout learning

We first analyzed the population-level activity of dSPNs and iSPNs across days, as previous reports have described dynamic changes in average neuronal activity in the DLS as animals learn new motor skills (Barnes et al., 2005; Yin et al., 2009; Lemke et al., 2019). Consistent with the notion that both the direct and indirect pathways of the basal ganglia contribute to motion (Cui et al., 2013; Klaus et al., 2017), we observed that the vast majority of dSPNs (74.9%) and iSPNs (79.2%) were positively modulated during locomotion (Figure 1E-H).

Similar proportions of dSPNs and iSPNs were positively modulated during bouts of locomotion on the first session (Figure 1E,F). However, across days the relative proportion of positively modulated dSPNs remained stable (Figure 1E), whereas the proportion of positively modulated iSPNs progressively decreased (Figure 1F). When we quantified the relationship between locomotion speed and Ca^2+^ event rates, we once again observed that dSPN activation was grossly similar between early (day 1 + day 2) and late (day 7 + day 8) stages of learning (Figure 1G), whereas iSPN activation significantly decreased by the late stage (Figure 1H).

Although the proportion of dSPNs being positively modulated remained stable across days, we asked if the magnitude of their activity during locomotion changed. We observed a significant decrease across days in the magnitude of locomotion-driven Ca^2+^ activation of both dSPNs and iSPNs (quantified as both dF/F and Ca^2+^ event rate; Figure 1I,J). Restricting our analysis to the beginning (onsets) or the end (offsets) of motion bouts yielded similar results, as both dSPNs and iSPNs exhibited decreased activation during both onsets and offsets of motion (Figure 1K-N). Together, these results show that population-level neuronal activation in the dorsal striatum in both the direct and indirect pathways progressively decreases as animals learn to self-generate locomotion on the running wheel.

### Action initiation recruits fewer SPNs across consecutive bouts of motion

With the observed across-day changes in SPN recruitment, we next asked if we could observe dynamic changes to SPN activity across consecutive bouts of locomotion within sessions belonging to the same day of training. When we quantified the proportion of dSPNs and iSPNs active during each of the first 8 consecutive locomotion bouts (Figure S3), we observed that dSPN (Figure S3A,B) and iSPN (Figure S3E,F) activation decreased across consecutive bouts of action initiation. This effect was not observed when looking at the first 8 consecutive offsets in either dSPNs (Figure S3C,D) or iSPNs (Figure S3G,H). Together, these results suggest that initial bouts of action initiation recruit larger populations of dSPNs and iSPNs compared to later bouts, where fewer striatal cells are engaged, even within the same session.

### Decreased activation of movement-nonspecific SPNs

Striatal neurons have been shown to be functionally heterogeneous, with individual SPNs displaying activity tuned to precise action motifs, temporal features, and task structures (Isomura et al., 2013; Rueda-Orozco and Robbe, 2015; Markowitz et al., 2018; Dhawale et al., 2021). We next asked if the significant decrease in striatal activity during learned locomotion was due to a global reduction in SPN activity, or if a decrease in a specific SPN population activity drove the effect. To this aim, we categorized dSPNs and iSPNs by the period of motion in which they were significantly active across a given session (Onset-only, Offset-only, Both, or Never).

Categorizing dSPNs and iSPNs into these functional classes revealed that despite the decreased number of SPNs recruited during locomotion, the average activity profiles of the remaining active SPN types were grossly unchanged (Figure 2A,B). When we compared the proportion of functional types across early and late stages of learning, we observed that the number of Onset-only and Offset-only SPNs remained stable across days (Figure 2C-F). However, the number of dSPNs and iSPNs that activated during both motion onset and offset (i.e., Both) significantly decreased across sessions (Figure 2C-H). Consequently, this decrease in Both-SPNs was mirrored by an increase in the proportion of SPNs that did not activate during motion onset or offset (i.e., Never; Figure 2E-H). Together, these results suggest that the decreased striatal activity across learning is specific to the population of dSPNs and iSPNs that grossly encode motion, leaving intact populations that are more specific to a precise action type, such as initiating or terminating locomotion.

**Figure 2.**
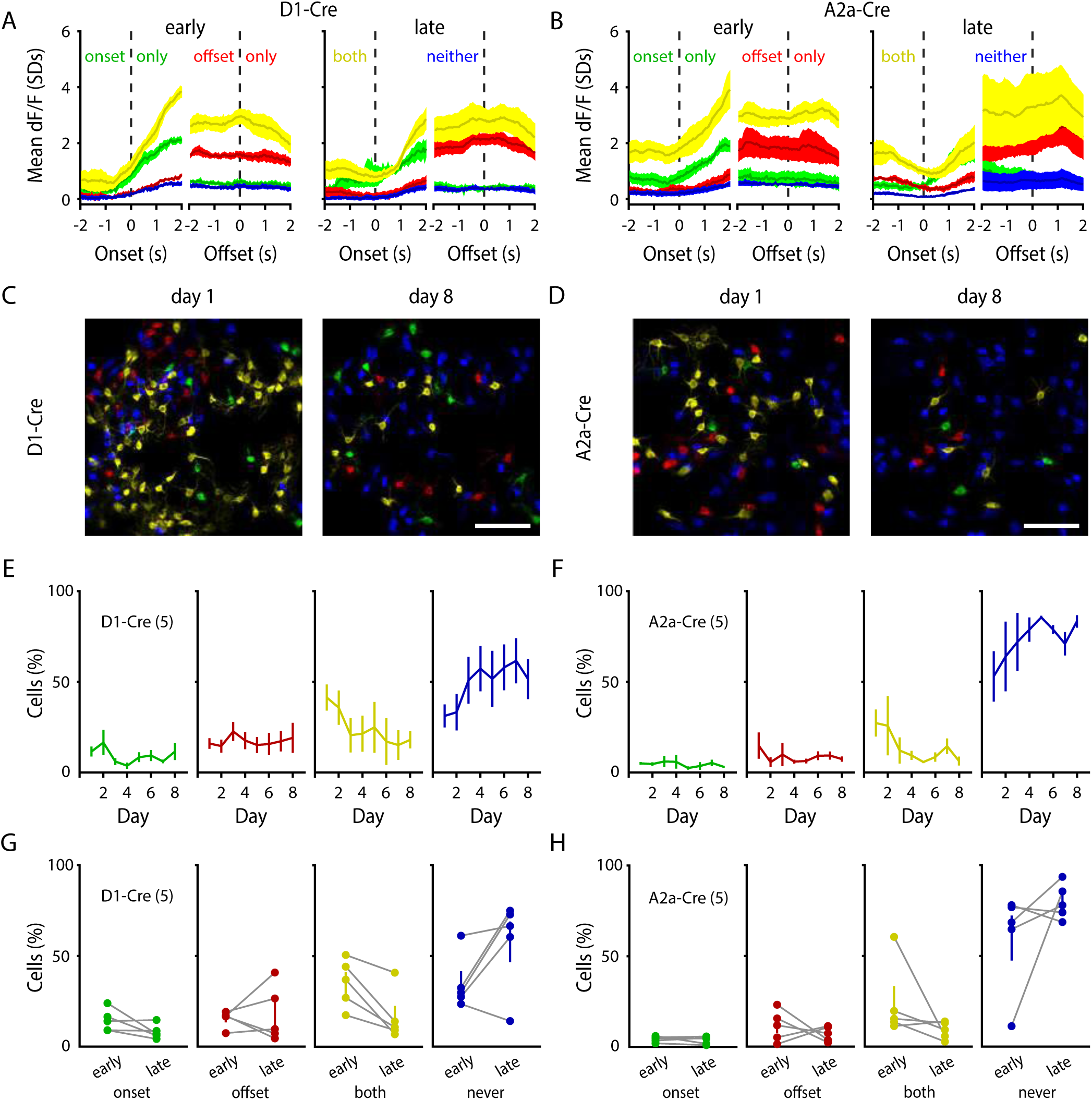
Specific decrease in the proportion of movement-nonspecific SPNs with learning. **A, B,** Average activity (standardized dF/F) of dSPNs (**A**) and iSPNs (**B**) where SPNs are categorized as being active only during motion onsets (‘onset only’, green), offsets (‘offset only’, red), active during both onsets and offsets (‘both’, yellow), or not active during either onsets or offsets (‘neither’, blue). Left- and right-side halves of panels depict average SPN activity during ‘early’ or ‘late’ sessions, respectively. **C, D,** Representative images of SPN locomotion activation types during session 1 (left) or session 8 (right). Scale bars: 100µm. **E,** Percentage of dSPNs across days that belong to the categories of ‘onset only’ (green; n = 5 mice / 8 days / 296 bouts; p = 0.462), ‘offset only’ (red; n = 5 mice / 8 days / 296 bouts; p = 0.167), ‘both’ (yellow; n = 5 mice / 8 days / 296 bouts; p = 1.905e-08), and ‘neither’ (blue; n = 5 mice / 8 days / 296 bouts; p = 2.605e-05). **F,** Percentage of iSPNs across days that belong to the categories of ‘onset only’ (green; n = 5 mice / 8 days / 318 bouts; p = 0.233), ‘offset only’ (red; n = 5 mice / 8 days / 318 bouts; p = 0.008), ‘both’ (yellow; n = 5 mice / 8 days / 318 bouts; p = 2.134e-05), and ‘neither’ (blue; n = 5 mice / 8 days / 318 bouts; p = 3.286e-08). **G, H,** Average activation types for ‘early’ (days 1+2) vs. ‘late’ (days 7+8) periods of learning. Data are mean ± SEM. Statistical significance was assessed by linear mixed-effects modeling (**E, F**).

### SPN motion-specific ensembles are formed from early active populations

Motor memories are thought to be encoded by precise ensembles of neurons (Laubach et al., 2000; Peters and Komiyama, 2017; Sheng et al., 2019; Josselyn and Tonegawa, 2020; Yuste and Yaksi, 2024). We observed that the number of onset-specific and offset-specific dSPNs and iSPNs remained stable over days while movement-nonspecific activity decreased. This raised an important conceptual question – are the identities of active SPNs stable across days of learning? To address this, we first performed a fate analysis where we searched for subsets of movement-activated SPNs during early training that persisted into late training days (as non-specific neurons generally became disengaged). Indeed, when quantifying the percentage of late training ensembles that were previously active during early training, we found that late training dSPN onset and offset ensembles were also significantly motion-activated at early training days (Figure 3A,B). Conversely, although early and late training iSPN ensembles overlapped for onsets (Figure 3C), we found that did not significantly overlap for offsets (Figure 3D).

**Figure 3.**
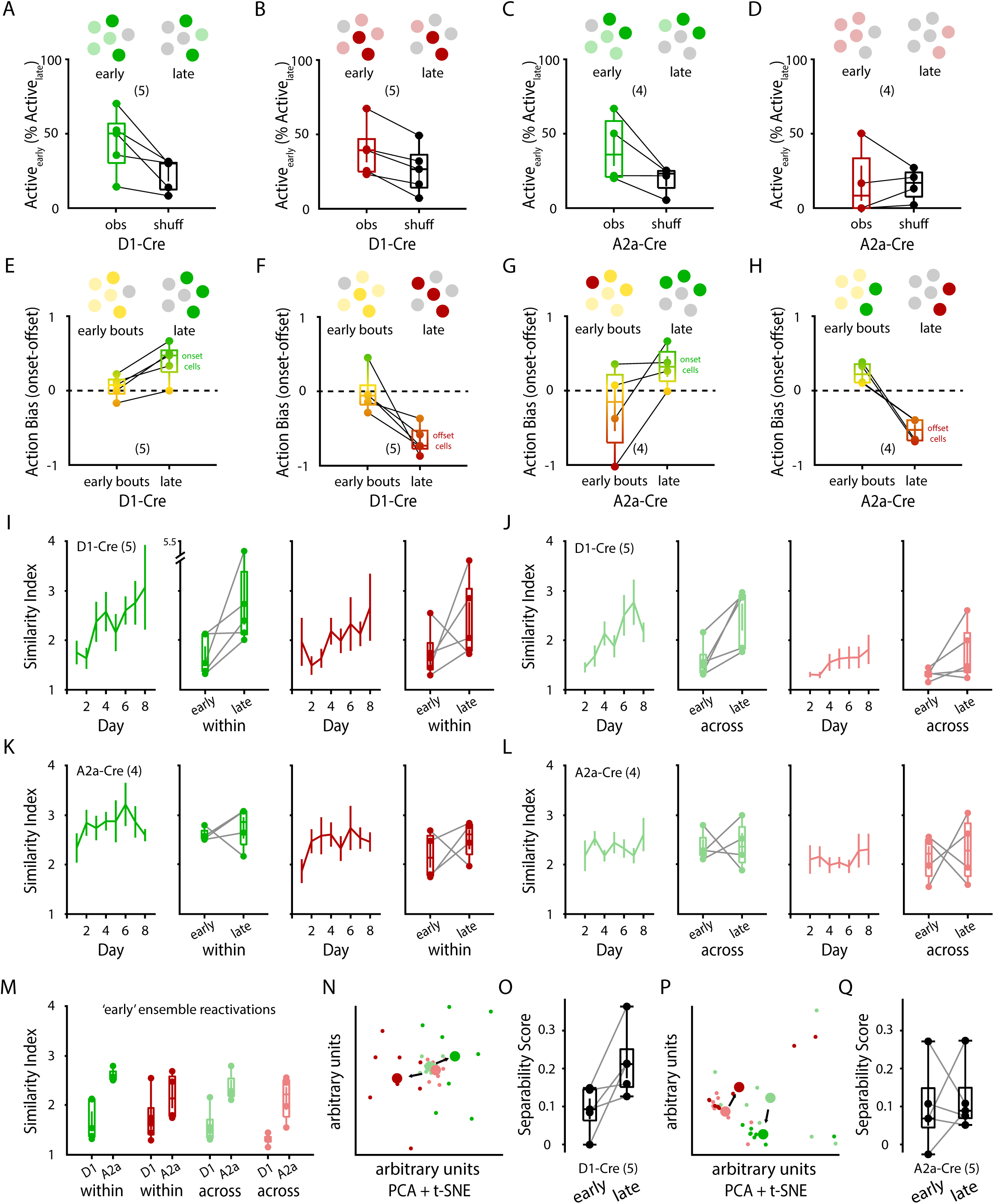
Refinement of movement-specific SPN ensembles throughout learning. **A, B,** Percentage of ‘late’ stage (days 7+8) onset (**A**) and offset (**B**) dSPN ensembles that were significantly active during ‘early’ stages (days 1+2). Left: observed activation overlap (obs), right: average overlap of 1000 shuffled activation vectors (shuff, see Methods) (n = 5 animals; dSPN onset: p = 0.042; dSPN offset: p = 0.006). **C, D,** Percentage of ‘late’ stage (days 7+8) onset (**C**) and offset (**D**) iSPN ensembles that were significantly active during ‘early’ stages (days 1+2). Left: observed activation overlap (obs), right: average overlap of 1000 shuffled activation vectors (shuff, see Methods) (n = 4 animals; iSPN onset: p = 0.104; iSPN offset: p = 0.927). **E, F,** Early vs. late Activation Bias averaged for each animal (across all dSPNs) for onset-specific (**E**; n = 5 animals; p = 0.007) and offset-specific (**F**; n = 5 animals; p = 0.026) dSPN ensembles. **G, H,** Early vs. late Activation Bias averaged for each animal (across all iSPNs) for onset-specific (**G**; n = 4 animals; p = 0.125) and offset-specific (**H**; n = 4 animals; p = 0.015) iSPN ensembles. **I,** Within-day Similarity Indices for onset-specific dSPN ensembles (green; n = 5 mice / 8 days / 381 onsets; p = 0.048) and offset-specific dSPN ensembles (red; n = 5 mice / 8 days / 401 offsets; p = 0.013) comparing bouts within days. **J,** Across-day Similarity Indices for onset-specific dSPN ensembles (green; 5 mice / 8 days / 174 onsets; p = 0.008) and offset-specific dSPN ensembles (red; n = 5 mice / 8 days / 174 offsets; p = 0.049) comparing bouts across days. **K,** Within-day Similarity Indices for onset-specific iSPN ensembles (green; n = 4 mice / 8 days / 379 onsets; p = 0.128) and offset-specific iSPN ensembles (red; n = 4 mice / 8 days / 373 offsets; p = 0.006) comparing bouts within days. **L,** Across-day Similarity Indices for onset-specific iSPN ensembles (green; 4 mice / 8 days / 140 onsets; p = 0.190) and offset-specific iSPN ensembles (red; n = 4 mice / 8 days / 140 offsets; p = 0.752). **I-L,** Right: box plots depicting summary of ‘early’ (days 1+2) vs. ‘late’ (days 7+8) average (per animal) Similarity Indices. **M,** Comparison of ‘early’ Similarity Indices between SPN types (‘within’ and ‘across’ days) (onset ‘within’ p = 0.004; offset ‘within’ p = 0.218; onset ‘across’ p = 0.009; offset ‘across’ p = 0.006). **N,** Representative PCA + tSNE dimensionality reduction of an animal’s dSPN population Ca^2+^ activity across bouts of onsets (green) and offsets (red), with ‘early’ (days 1+2) bouts depicted in lighter colors and ‘later’ (days 7+8) bouts depicted in darker colors. **O,** Separability Score (onsets vs. offsets) of all D1-Cre animals comparing the ‘early’ to ‘late’ stage dSPN population separability of actions (n = 5 mice; p = 0.046). **P,** Representative PCA + tSNE dimensionality reduction of an animal’s iSPN population Ca^2+^ activity across bouts of onsets (green) and offsets (red), with ‘early’ (days 1+2) bouts depicted in lighter colors and ‘later’ (days 7+8) bouts depicted in darker colors. **Q,** Separability Score (onsets vs. offsets) of all A2a-Cre animals comparing the ‘early’ to ‘late’ stage iSPN population separability of actions (n = 5 mice; p = 0.255). Data are mean ± SEM. Box plots are depicted as mean (center), first/third quartile (lower/upper box limits), and minima/maxima (bottom/top whiskers). Statistical significance was assessed by two-sided paired t-tests (**A-H, O, Q**), linear mixed-effects modeling (**I-L**), and two-sided unpaired t-tests (**M**).

We next asked what the activation type (i.e., onset, offset, or both) of the eventual ensembles (day 7 + day 8) were during the early sessions (day 1 + 2). By quantifying the ‘Action Bias’ of these SPNs as how activated they were during early onsets vs early offsets, we found that the eventual dSPN onset and offset ensembles were activated early on during both actions (Figure 3E,F). In contrast, eventual iSPN onset ensembles had an early slight offset-bias (Figure 3G) and eventual iSPN offset ensembles had significant early onset-bias in activation (Figure 3H). These results suggest that dSPN ensembles are formed from early action nonspecific dSPNs while iSPNs ensembles are formed from a mix of action nonspecific iSPNs and iSPNs that show a very early bias to the opposing action.

### Movement-specific SPN ensembles are refined throughout learning

To further characterize the stability of movement-specific SPN ensembles throughout learning, we measured the similarity of SPN neurons reactivated across bouts of onsets and offsets. Strikingly, we observed that the bout-to-bout dSPN populations activated during motion onsets or offsets significantly increased in similarity throughout days of learning (Figure 3I). This effect held true when comparing the similarity scores of dSPN onset/offset ensembles within the same day or of ensembles across adjacent days (Figure 3J). These results suggest that the direct pathway progressively refines stable populations of dSPNs that are to be activated during specific types of actions.

Surprisingly, iSPN populations did not undergo a significant increase in within-day or across-day similarities during onsets or offsets (Figure 3K,L). Instead, iSPN similarities during the first training days already started significantly higher than those of dSPNs (Figure 3M). Thus, whereas the direct pathway progressively decreases its activation over days while simultaneously refining motion-specific dSPN ensembles, the indirect pathway also decreases its activation over days, but the active iSPNs already start off more consistently reactivated across bouts of motion. Consistent with this notion, when we used dimensionality reduction to compare ensemble activations between ‘early’ and ‘late’ bouts of motion, we found that low-dimensional dSPN encodings of onsets and offsets became progressively more separable (Figure 3N,O), whereas iSPN encodings did not (Figure 3P,Q). Together, these results suggest that the dorsal striatum refines populations of striatal SPNs engaged during bouts of motion onset or offset.

### Striatal movement-specific activity becomes more efficient throughout learning

We next asked if these dynamic changes in dSPN and iSPN activity resulted in changes in population-level encoding of movements in the striatum. To quantify the relationship between population SPN activity and their encoding of self-generated locomotion, we trained linear regression models (see Methods) to predict running velocity from SPN activity. To distinguish between onset and offset encodings, we utilized separate models for motion onsets or motion offsets. When using all imaged dSPNs or iSPNs in the models, we found that model performance did not change throughout days (Figure 4A-D), suggesting that overall information encodings in the striatum were unchanged throughout learning. Because we observed that activated SPN ensembles became more similar across training days (Figure 3), we hypothesized that late training days might engage ensembles that are more predictive of motion. However, when quantifying model performance trained on discrete subsets of SPNs, we found that model performance was not significantly changed across days, even when only including the ‘best’ 2-8 SPNs (Figure. 4E-H; see Methods). These results are consistent with our observations that motion-specific dSPN and iSPN ensembles are refined from populations of SPNs active during initial learning (Figure 3A-H), rather than emerging as a newly activated population throughout learning.

**Figure 4.**
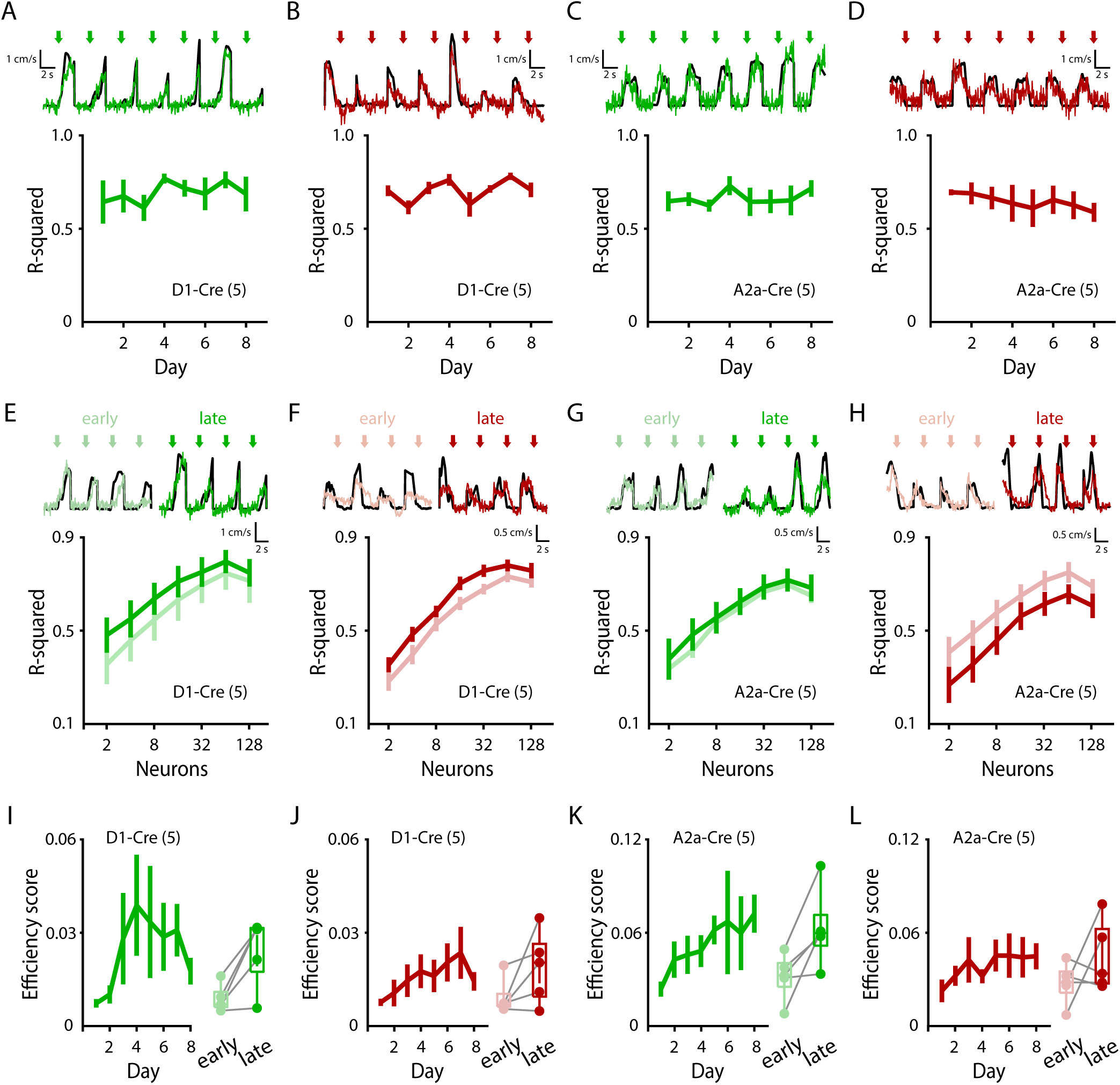
Striatal movement-specific activity becomes more efficient throughout learning. **A, B,** Average predictive performance across days of action-specific linear regression models fitting dSPN Ca^2+^ activity to animal velocity, trained on subsets of onsets (**A**; n = 5 mice; p = 0.420) or offsets (**B**; n = 5 mice; p = 0.142). **C, D,** Average predictive performance across days of action-specific linear regression models fitting iSPN Ca^2+^ activity to animal velocity, trained on subsets of onsets (**C**; n = 5 mice; p = 0.470) or offsets (**D**; n = 5 mice; p = 0.544). **E, F,** Model performance of action-specific (onset-specific, **E**; offset-specific, **F**) linear regression models trained on subsets of dSPNs of varying sizes (see Methods; onsets: n = 5 mice, p = 0.474; offsets: p = 0.982). **G, H,** Model performance of action-specific (onset-specific, **G**; offset-specific, **H**) linear regression models trained on subsets of iSPNs of varying sizes (see Methods; onsets: n = 5 mice, p = 0.841; offsets: p = 0.337). **I, J,** Average ‘Efficiency Score’ (see Methods) plotted for dSPN encodings of onsets (**I**; n = 5 mice; p = 0.017) or offsets (**J**; n = 5 mice; p = 0.074). **K, L,** Average ‘Efficiency Score’ (see Methods) plotted for iSPN encodings of onsets (**K**; n = 5 mice; p = 0.049) or offsets (**L**; n = 5 mice; p = 0.128). **I-L,** Right: comparison of ‘early’ (days 1+2) vs. ‘late’ (days 7+8) stage ‘Efficiency Scores’. Data are mean ± SEM. Box plots are depicted as mean (center), first/third quartile (lower/upper box limits), and minima/maxima (bottom/top whiskers). Statistical significance was assessed by repeated measures 1-way ANOVA with multiple comparisons (**A-D**), 2-way repeated measured ANOVA with multiple comparisons (**E-H**), and two-sided paired t-tests (**I-L**).

Finally, as velocity prediction performance was unchanged throughout learning, but the overall numbers of dSPNs and iSPNs active at onsets and offsets progressively decreased, we calculated the relative efficiency of striatal encoding of motion (quantified as R^2^ / #SPNs). Indeed, we found that encoding efficiency of motion onsets significantly increased throughout training days for dSPNs and iSPNs (with offset encodings following the same trend but not reaching significance) (Figure 4I-L). These results suggest that as dSPN and iSPN activity is refined throughout learning and motion-nonspecific activity decreases, the striatum becomes more efficient and consistent at encoding specific movements.

## DISCUSSION

In this study, we aimed to characterize how striatal SPN activity encodes (i.e., predicts) distinct actions and how these encodings evolve throughout motor learning. In combination, our findings suggest that motion-specific dSPN ensembles emerge from motion-nonspecific dSPNs, which are progressively deactivated or refined throughout learning. Although iSPN ensembles did not meet significance criteria for longitudinal reactivation (Figure 3C,D), the high similarity of iSPN ensemble activation across movement bouts (Figure 3K,L), and progressive deactivation of movement-nonspecific iSPNs (Figure 2F,H) suggest that iSPN ensembles are also precisely defined and dynamic with learning, although perhaps to a lesser degree than dSPNs. At the population level, overall striatal activity decreases while refining and preserving these movement-specific ensembles, resulting in a more efficient encoding of movement behavior. Our longitudinal results provide a link between early- and late-learning SPN dynamics to elucidate how striatal ensembles are formed during learning.

Decades of research have yielded a current model of the basal ganglia wherein dSPNs and iSPNs exert largely distinct and at times opponent commands over actions (Gittis and Kreitzer, 2012; Calabresi et al., 2014), but with the precise recruitment of both pathways being required for normal action selection, control, and learning (Cui et al., 2013; Klaus et al., 2017; Arber and Costa, 2022). Our observations are consistent with a model involving the activation of both dSPNs and iSPNs across distinct (even opposing) actions, as both locomotion onsets and offsets recruit both pathways (Figure 1). Our data support a role for the direct pathway in promoting specific actions, as we observe during early learning that it is the dSPNs that show some activation to the performed action that are progressively reactivated and refined throughout learning (Figure 3A-F) and are consistently recruited across bouts during later sessions (Figure 3I,J). Our data are also consistent with a role for precise, concurrent indirect pathway activation during movement, as the iSPN activity being recruited to both onsets and offsets exhibits high reactivation similarity across bouts and days (Figure 3K-M). A major difference between dSPNs and iSPNs in our data is that iSPN action-specific ensembles show an early bias for the opposing action (e.g., offset iSPNs showing an early activation bias for onsets despite later being offset-specific; Figure 3H). One possible interpretation is that iSPN ensembles are initially reinforced to the action context opposite to their eventual recruitment type (Lindsey et al., 2024). This would be consistent with the role of the indirect pathway in precisely inhibiting actions in opposition to the learned action. For example, early onset bouts may form iSPN ensembles that are then later activated during offset events in order to indirectly promote offsets by inhibiting onsets (and vice versa) (Freeze et al., 2013; Tecuapetla et al., 2016; Lindsey et al., 2024). In addition to the observed ‘type reversal’, our observations that iSPNs undergo a more dramatic change in their motion-modulation (Figure 1F,H) are also consistent with this.

For both dSPNs and iSPNs we observe that early sessions recruit significantly more activated SPNs than later sessions (Figure 1K-N), an effect that can also be observed on a trial-to-trial basis, as initial bouts within a session recruit more dSPNs and iSPNs than later ones (Figure S3). It is possible that the gross initial activation of striatal cells reflects an expansive exploration of neuronal space throughout the DLS, allowing for the sampling and recruitment of the eventual ensemble cells to be refined (Vicente et al., 2016; Lee and Sabatini, 2021; Kondapavulur et al., 2022). Alternatively, the greater activation may simply reflect a greater excitatory tone onto SPNs from the cortex and thalamus as the cortico-basal ganglia activity is progressively refined as animals learn to smoothly locomote (Tang et al., 2007; Hwang et al., 2019; Cataldi et al., 2022).

Learning has been shown to coincide with strengthening activity between the cortex and the striatum at the cellular and synaptic levels (Pisani et al., 2005; Koralek et al., 2012; Athalye et al., 2018; Lemke et al., 2019; Lemke et al., 2021). The formation of cortical ensembles has been mechanistically linked to the dynamic reorganization of synaptic inputs onto cortical cells during learning (Peters and Komiyama, 2017; Roth et al., 2020; Albarran et al., 2021), but a role for these cortical ensembles in instructing downstream learning in the striatum has been challenging to describe mechanistically. Because the cortico-basal ganglia are thought to be organized into functional loops, with striatal outputs ultimately feeding back on “upstream” cortical activity through basal ganglia outputs and corticothalamic activity, it has been difficult to mechanistically isolate plasticity within a particular link in this chain.

Recent *ex vivo* findings suggest that cortical ensembles directly drive synaptic plasticity changes onto downstream SPNs, providing a possible mechanism linking cortical and striatal ensembles during learning (Hwang et al., 2022; Pimentel-Farfan et al., 2022). Indeed, it is possible that the SPN ensemble refinement we observed in this study is part of a broader functional refinement reverberating simultaneously throughout the cortico-basal ganglia circuitry, driven by synaptic plasticity mechanisms at each point. Our observed longitudinal striatal changes are consistent with this model, as the time course of nonspecific SPN deactivation and ensemble refinement across days and within days (i.e., across bouts) is consistent with the time courses from previous studies using motor skill learning tasks (Yin et al., 2009; Peters and Komiyama, 2017; Lemke et al., 2019; Albarran et al., 2021). Future experiments simultaneously measuring cortical and striatal ensemble formation during learning will be necessary to elucidate a detailed mechanism for dynamic information flow during learning.

The striatum is critical to various motor and cognitive processes, reflected in its wide-ranging mixed and multimodal inputs combined from across the cortex, thalamus, hippocampus, amygdala, and the midbrain. As a functional cornerstone to such diverse processes, the striatum needs to translate these wide-ranging convergent inputs into stable and efficient encodings. Indeed, the striatum must balance the flexibility required to integrate these broad inputs into SPN ensembles, while simultaneously maintaining a robust stability of these memory substrates across distinct contexts (Mau et al., 2020; Pérez-Ortega et al., 2021). As SPNs possess thousands of dendritic spines but receive few inputs from each presynaptic partner (Wilson, 2004), they are well poised to serve as an integrative substrate from which diverse ensembles are generated and reinforced through plasticity mechanisms (Calabresi et al., 2007).

Our data supports the notion that learning new actions involves the initial recruitment of a broad array of SPNs from which a precise ensemble is refined over repeated trials and sessions. The initial burst of SPN activation we observe in early days (and early bouts within the same day) may reflect an intrinsic encoding flexibility that allows the striatum to generate new ensembles from pools of motion-nonspecific SPNs (Dhawale et al., 2017; Hwang et al., 2019; Lee and Sabatini, 2021). Indeed, reinforcement through synaptic plasticity rules likely underlies the SPN ensemble refinement we observe, although future experiments are needed that manipulate neuromodulators such as dopamine, acetylcholine, the endocannabinoid system, and other mechanisms of striatal synaptic plasticity while longitudinally recording striatal ensembles (Maltese et al., 2021; Ma et al., 2022; Krok et al., 2023; Albarran et al., 2023; Yang et al., 2023).

In this light, encoding flexibility and encoding redundancy go hand in hand, as redundancy in the neural code is documented throughout the nervous system, particularly in sensorimotor regions (Narayanan et al., 2005; Butz et al., 2007; Nigam et al., 2019). Some redundancy in neuronal encoding is believed to be advantageous, such as increasing signal-to-noise (i.e., robustness) or protection from misfires and degradation (Rolls and Treves, 2011; Raman et al., 2019). Indeed, although onsets and offsets in theory only need to be encoded by a single SPN each, encoding through SPN ensembles (i.e., our observed ∼15% of imaged SPNs) is a redundancy that may be advantageous to drive the behavior more robustly from bout to bout. Of course, the cost of redundancy is efficiency, as utilizing the fewest number of resources (e.g., SPNs) to encode a given action is desirable both energetically and from an information storage standpoint to minimize conflicting commands and increase signal-to-noise (Cai et al., 2016; Denève and Machens, 2016; Tang et al., 2019). To this aim, the striatal dynamics we observe in this study are in line with a process that balances redundancy with efficiency, as initial learning coincides with the recruitment of a vast, nonspecific, and largely redundant population of SPNs that are then refined into smaller, specific ensembles that yield more efficient encodings of actions.

## METHODS

### Mice

All experiments were performed in accordance with protocols approved by the Stanford University Animal Care and Use Committee in keeping with the National Institutes of Health’s *Guide for the Care and Use of Laboratory Animals*. Animals were kept at a 12hr:12hr light/dark cycle at a room temperature of 22°C with humidity control (30-70%). For our experiments, we used heterozygous Drd1a-Cre and A2a-Cre mice (Gong et al., 2007). We used both male and female mice, aged ∼8 weeks at the start of behavioral experiments.

### Surgical procedures

We performed surgeries on animals under isoflurane anesthesia (1.5-2.5% in 0.5L/min of O_2_). To drive the expression of GCaMP6f in the striatum of either A2a or Drd1-Cre mice, we stereotaxically injected 400nl of AAV1-syn-Flex-GCaMP6f-SV40 (100833-AAV1, Addgene) into the dorsolateral striatum (from bregma, anterioposterior: 1.0mm; mediolateral: 2.25mm; and from dura, dorsoventral: -2.45). Viral construct was injected using a pulled glass pipette at a rate of 40nl/min. After the injection, we waited for 10 min before withdrawing the pipette. We then sutured the incision and let animals recover in their cages overnight before a second surgery where we implanted an optical guide tube for GRIN lenses.

We fabricated the GRIN lenses guide tube by gluing (UV curable glue Norland #81) a 2mm-diameter #0 glass coverslip to the tip of a 3.8mm long 18G stainless steel tube (1.27mm OD, 1.06mm ID, McMaster-Carr). Excess glass was removed using a polishing wheel (Ultratec basic polisher, 30mic silicon abrasive disc).

For the implantation of GRIN lens guide tubes, we anesthetized previously injected animals with isoflurane (1.5-2% in 0.5L/min of O_2_). Using the same coordinates where the glass pipet was inserted, we generated a new craniotomy using a 1.4 mm diameter drill bit (F.S.T.). With a 27G blunt needle we aspirated the cortex until the striatum was visible (dorsoventral:-2.0mm from dura). Then the guide tube was lowered at -2.35mm from dura. After insertion we sealed the gap between the cannula and the skull using n-butyl cyanoacrylate adhesive (Vetbond, 3M), before covering the area with dental cement (Metabon, Parkell). Finally, a second batch of dental cement was used to attach an aluminum head bar to the cranium. We allowed the animals to recover and express GCaMP6f for at least 3-4 weeks before behavioral/imaging experiments.

### Behavioral apparatus and training

Our training device consists of a 14cm diameter microfoam wheel connected to a shaft encoder (MA3-A10-250-N, US Digital); the encoder provides a readout of the wheel revolutions and hence the travel distance and speed. Using custom code controlled by an Arduino board, we set a random reward reward delivery that activates a peristaltic silicon pump (Adafruit), delivering 0.2ml of water through a blunt needle at random intervals not shorter than 8 seconds. The blunt needle is connected to a capacitive touch sensor (AT42QT1010, Sparkfun) that records every time animals lick to consume the water rewards (lick-o-meter). We recorded the encoder, pump activation, and lick-o-meter signals along with the timing pulses from the microscope’s frame clock using custom made codes with a digital data acquisition system (LabView,NI). Mice underwent a water-restriction protocol whereby daily water intake was limited to ∼1mL daily. We monitored the animal’s weight daily to avoid more than a 20% drop from their starting weight and provided supplementary water as necessary. Animals training consisted of exposure to the running wheel for 8 consecutive days for ∼20 min. Mice were head-restricted with their limbs resting on the wheel, and were allowed to run at will. During this training their neuronal calcium activity was recorded using two-photon imaging.

### Two-photon microscopy

We used a 25 x /1.0 NA water immersion objective lens (Olympus) for striatal imaging, covering a 250 × 250 μm field-of-view (FOV) through a 1 mm diameter GRIN probe #1050-002176, Inscopix). We carefully returned to the same FOV every imaging session during the 8 days of training based on the vasculature and neuronal population patters (Figure S1). A Ti:Sapphire laser (MaiTai Deepsea, Spectra-Physics) provided 920 nm excitation for two-photon imaging. We used 60-80mW laser power through the objective, as measured above the implanted GRIN probe. We detected fluorescence signals using a GaAsP photomultiplier tube (H10770PA-40, Hamamatsu) and band-pass fluorescence emission filters (FF01-520/60 for GCaMP, FF01-590/36 for tdTomato; Semrock) adapted to a custom-built two-photon microscope system with a resonant scanner (LotosScan, Suzhou Institute of Biomedical Engineering and Technology) and a 25 x / 1.0 NA water immersion objective lens (Olympus).

All videos were acquired at 512 × 512 pixel resolution at ∼25 Hz frame rate. The precise temporal alignment between the Ca^2+^ imaging videos and the behavior was established by triggering the recording of all components at the beginning of each session using LotosScan.

### Image processing and extraction of calcium signals

All recorded two-photon videos underwent the same pre-processing pipeline. First, we corrected for any DC offset in the pixel values. For each frame, we computed the minimum pixel value over the entire image. We then averaged this value over all frames and subtracted the result from all pixels in the movie. Next, we corrected for brain motion using piecewise rigid motion correction (NoRMCorre) (Pnevmatikakis and Giovannucci, 2017) using 32 X 32 pixel patches of each image. We then corrected for slow changes in FOV brightness throughout recordings (e.g. caused by mild bleaching of the calcium indicators). We did this by calculating the mean pixel value of the first image, fitting an exponential curve (a∗exp(−b∗t)+c) to the brightness as a function of each frame, and dividing each frame by its fitted brightness. Finally, we z-scored each pixel using its mean value and standard deviations over all frames.

To identify cells and extract their Ca^2+^ activity traces from the z-scored Ca^2+^ videos, we used constrained non-negative matrix factorization (CNMF) (Pnevmatikakis et al., 2016) or EXTRACT (H. Inan et al., 2021) cell-sorting algorithms. Candidate neuron traces were manually inspected to confirm the quality of their segmentation (spatial filters) and signal-to-noise ratio. Neurons that failed to satisfy both of these attributes were excluded from future analysis.

After extracting spatial filters corresponding to individual neurons (using CNMF or EXTRACT), we truncated the filters by setting to zero the weights of all pixels with values less than 1.5 s.d. above the mean pixel weight, averaged across the entire spatial filter. We applied these truncated spatial filters to the raw fluorescence videos to extract fluorescence traces for classified neurons. Because our data consisted of eight consecutive recording sessions, we inspected if any classified neurons of each day were present or not in other days (Figure S4). This allowed us to resolve if a cell was not detected on a given day due to a failure of the cell-sorting algorithms, or if the cell exhibited low Ca^2+^ activity. Thus, for each pair of days, we took the original spatial filters generated by the cell-sorting algorithms and computed cross-day cell matchings. Filters without any matches were seeded across days, and the Ca^2+^ activity of these candidate filters were manually inspected for clear neuronal Ca^2+^. Finally, we merged the confirmed filters for a given day, removing duplicates by generating “union” filters. Fluorescence traces were then extracted by applying the “union” filters to the z-scored Ca^2+^ videos via least-squares.

### Detection of Ca^2+^ events and active cells

Timepoints of transient activation were calculated individually for each neuron. Standardized dF/F vectors for each neuron were smoothed using a moving average with a span of 0.5 s. After calculating the derivatives of these smoothed vectors, events were identified as the peaks in the activity that exceeded a threshold of 0.4 dSD/dt. Cells were classified as active if their standardized dF/F activity exceeded a threshold of 2 SD. Cells were further classified as Onset-only, Offset-only, of Both if they were active at onsets ± 2 s, offsets ± 2 s, or both at onsets and offsets, respectively.

### Classification of movement types

Motion onsets and offsets were independently determined using threshold criteria. Motion onsets were defined as timepoints preceded by a 2 s period where the rod velocity maintained below 0.09 cm/s and proceeded by a 2 s period where the rod velocity exceeded 0.35 cm/s at least once. Motion offsets were defined as timepoints preceded by a 2 s period where the rod velocity exceeded 0.30 cm/s at least once and proceeded by a 2 s period where the rod velocity maintained below 0.09 cm/s. In cases where such onsets or offsets were closer than 2 seconds apart, the later timepoint was omitted. For motion-activation analysis, timepoints were classified as motion (motion_ts) between every motion onset and the earliest proceeding timepoint where the average velocity in a 0.7 s window fell below 0.09 cm/s. Non-motion timepoints (nonmotion_ts) were defined as all timepoints that were not motion_ts.

### Quantifying the proportion of positively modulated cells

For each neuron, the number of active timepoints (active_ts) was calculated as the number of timepoints where its standardized dF/F was greater than 2 standard deviations. A neuron was classified as positively modulated if the proportion of active_ts / motion_ts was greater than the proportion of active_ts / nonmotion_ts. The final proportion of positively modulated cells was then determined to be the number of positively modulated cells divided by the total number of cells imaged on that day.

### Bout to bout activation analysis

To analyze the proportion of SPNs activated across consecutive bouts (e.g., onsets), we first extracted timepoints corresponding to the first consecutive 8 onsets (or offsets), and then categorized neurons for each onset by whether or not their standardized dF/F exceeded 3 SD within a 2 s window centered at each onset (or offset). We then averaged these proportion curves across animals (per genotype) and across days in order to quantify the average bout-to-bout activation of SPNs across the first 8 consecutive bouts of any session.

### Early-to-late ensemble reactivation fate analysis

In order to quantify the extent to which late-stage ensembles were already activated during early-stages, we compared activity-vectors across days for onsets or offsets. For each animal, we took the activation vectors for the first two days (early) or the last two days (late) and created ‘early’ and ‘late’ activation vectors by defining a neuron as active if it active on both of the two days. We then determined the amount of pre-existing activation by calculating the % of ‘late’ neurons that were also active during ‘early’ training.

### Early-to-late ensemble type fate analysis by quantifying Action Bias

To retroactively determine the activation ‘types’ that SPNs belonged to before they became eventual action-specific ensembles, we took the activation vectors for the last two days (late) and created ‘late’ activation vectors (averaged over bouts of both days) for onsets and offsets. Averaging over the early bouts (bout 1 and 2) of early days (days 1-3) we then assigned each SPN as ‘1’ if they were on average active during onsets, ‘-1’ if active during offsets, and ‘0’ if active during both onsets and offsets (else they were omitted), yielding an ‘early bouts’ vector that we then averaged for each animal to create an overall ‘Action Bias’ for each animal.

### Ensemble activity similarity index

To quantify the similarity between two neuron ensembles for bouts of onsets / offsets within the same day or across days, analysis was restricted only to neurons that were detectable across all 8 days of behavior. For each onset / offset bout, ensemble vectors were created such that the i^th^ value in the vector was 1 if the i^th^ neuron was active (see above) during that movement bout, otherwise the value was 0. In order to compare the similarity between two ensembles, we computed the Jaccard index: the number of shared activated neurons in the respective ensemble vectors, divided by the total number of activated neurons across both ensembles. Each Jaccard index was normalized to the average Jaccard index of 1000 shuffles of the two ensemble vectors.

### Action Separability Score

In order to quantify how separable onset and offset actions were at the level of population Ca^2+^ activity, we first generated a matrix N consisting of rows corresponding to each onset and offset on a given day, and the columns being horizontally concatenated timeseries of each SPN imaged on that day (where the timeseries are 4 second windows centered around the onset or offset corresponding to the row). This N matrix then underwent dimensionality reduction, first using PCA (preserving 8 principal components), and then using t-SNE using the Chebyshev distance metric. Separability scores were then calculated as the [day1 + day2] average or [day7 + day8] average silhouette cluster scores for ‘early’ and ‘late’ stages respectively.

### Peri-event regression analysis

To measure the ability of SPN Ca^2+^ activity to predict rod velocity during onsets and offsets, we first isolated 4-second windows around motion onsets or offsets, and temporally concatenated these bouts to create a singular timeseries vector of motion onsets or offsets for each day. Using the same timestamps, the Ca^2+^ activity of each SPN was concatenated to capture their activity before and after these events. We then fit linear regression models using the Ca^2+^ matrices to predict the rod onset or offset vectors. Models were k-fold cross-validated (where k was equal to the number of onsets or offsets). Using this method we used the adjusted R-squared metric to quantify the relationship between actions and population-level activity across days.

### N-neuron regression analysis

In order to measure the amount of information encoded by subsets of SPNs of increasingly larger sizes, we trained linear regression models as described above, but this time only utilizing 2, 4, 8, 16, 32, 64, or 128 neurons. To determine which neurons to use in the N-neuron models we used LASSO regression, varying the lambda parameter to achieve a model sparsity equal to the desired N count. Once the N neurons were identified, we then fit a new linear regression model using only the N neurons using the same cross-validation procedure described above.

### Efficiency score

To determine the efficiency of striatal encoding of actions, we used the per-neuron regression curves (see above) and the previously quantified number of active neurons per day and quantified the R-squared value corresponding to that number of neurons. Specifically, X curves were fitted to the per-neuron regression curves of each day, to match an R-squared value to a given number of activated neurons. This metric, which we the named efficiency score, reflects the amount of information encoded divided by the number of neurons activated to yield that R-squared value.

### Statistics

Animal subjects were randomized within experimental blocks to yield equal sampling of experimental conditions. All experiments were conducted in a blind fashion (i.e., experimenter did not know the mouse genotype). Data distribution was assumed to be normal, but this was not formally tested (hence individual data points are presented in all statistical comparisons). Repeated measurements (e.g., measure of average rod velocity across days) were analyzed using 1-way or 2-way repeated measures ANOVA with post-hoc tests. All two-sample comparisons (e.g., Similarity scores) were analyzed with t-tests. Unless otherwise specified, two-sided statistical tests were conducted and data is presented as mean ± SEM (standard error of the mean), with all statistical tests, statistical significance values, and sample sizes described in the figure legends.

## Acknowledgments

This study was supported by grants from the NINDS/NIH R01NS091144 (J.B.D.), GG gift fund (J.B.D.), Catalyst grant from the Phil & Penny Knight Initiative for Brain Resilience at the Wu Tsai Neurosciences Institute, Stanford University (J.B.D.), HHMI Gilliam Fellowship (E.A.), NSF GRFP (E.A.), and Stanford DARE Fellowship (E.A.). We thank Xiaobai Ren for technical support, Tony Hyun Kim for longitudinal calcium imaging and cross-day alignment consultation, Alex Williams and Vivek Athalye for computational methods discussion, and the members of Ding lab for valuable discussions.

## Author contributions

O.J., E.A., and J.B.D. conceptualized the project and designed the experiments. O.J. performed animal surgeries and behavioral experiments. Y.W. provided crucial scientific insight during early project stages. O.J., E.A., E.N.A., and J.B.D. analyzed the data. O.J., E.A., and J.B.D. wrote the manuscript with input from all authors.

---

**Figure S1.**
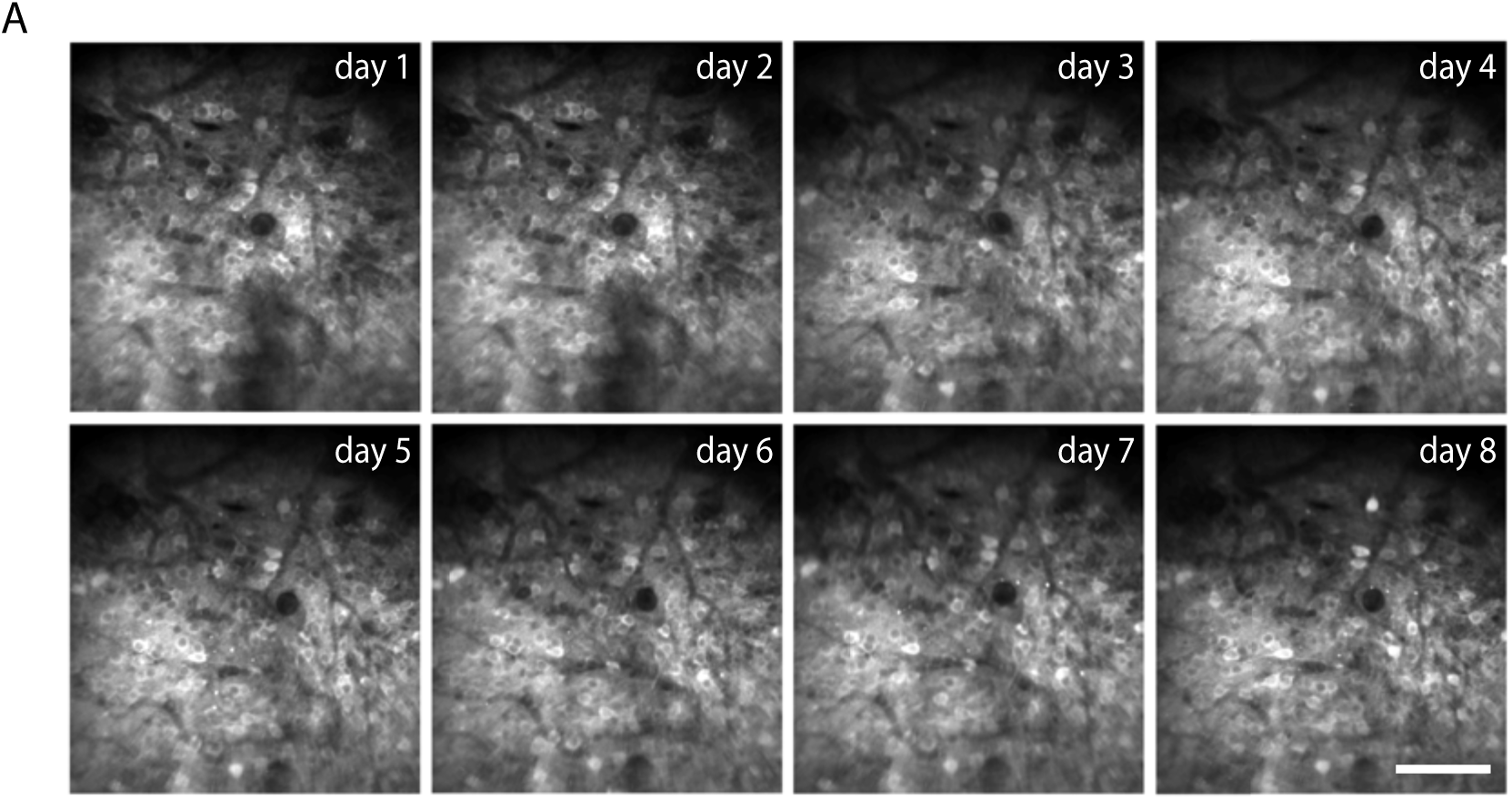
Longitudinal imaging of striatal SPNs. **A,** Representative imaging of the DLS through the 2-photon microscope, depicting consistent revisiting of the same field of view across sessions. Scalebar: 100 µm.

**Figure S2.**
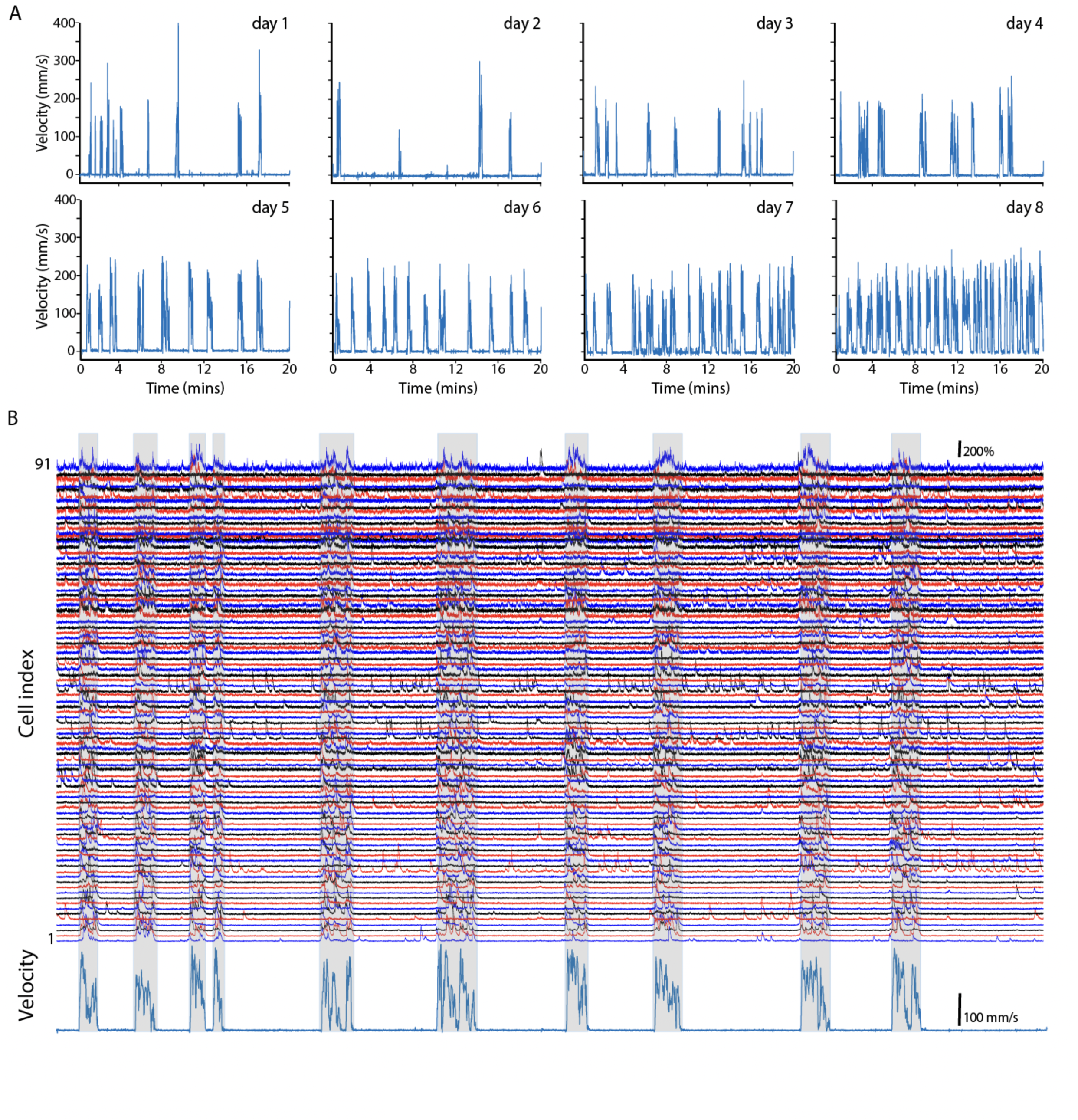
Imaging striatal SPNs throughout locomotion paradigm. **A,** Representative plots of an animal’s velocity across days. **B,** Representative Ca^2+^ traces (top) of imaged SPNs as mice perform self-generated bouts of locomotion (bottom).

**Figure S3.**
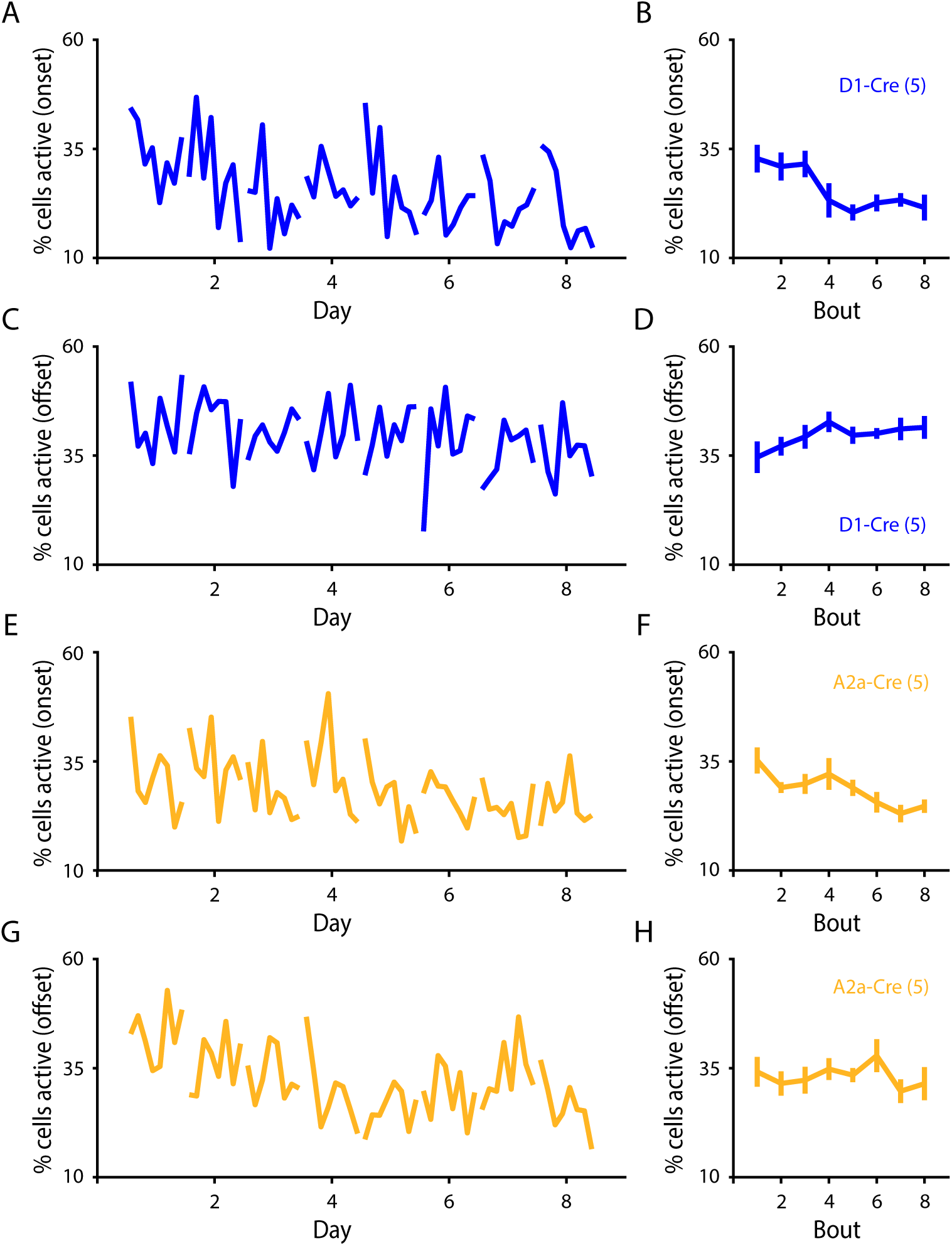
Bout-to-bout decrease in SPN activation during action initiation. **A,** Percentage of dSPNs active during the first 8 motion onsets of each day (averaged across animals). **B,** Activation percentages for the first 8 bouts averaged across days revealing a significant decrease in dSPN activation across consecutive motion onsets (n = 5 mice; p = 0.019). **C, D,** dSPN activation across the initial 8 bouts of motion offsets (n = 5 mice; p = 0.369). **E,** Percentage of iSPNs active during the first 8 motion onsets of each day (averaged across animals). **F,** Activation percentages for the first 8 bouts averaged across days revealing a significant decrease in iSPN activation across consecutive motion onsets (n = 5 mice; p = 0.014). **G, H,** iSPN activation across the initial 8 bouts of motion offsets (n = 5 mice; p = 0.352). Data are mean ± SEM. Statistical significance was assessed by repeated measures 1-way ANOVA with multiple comparisons (**B, D, F, H**).

**Figure S4.**
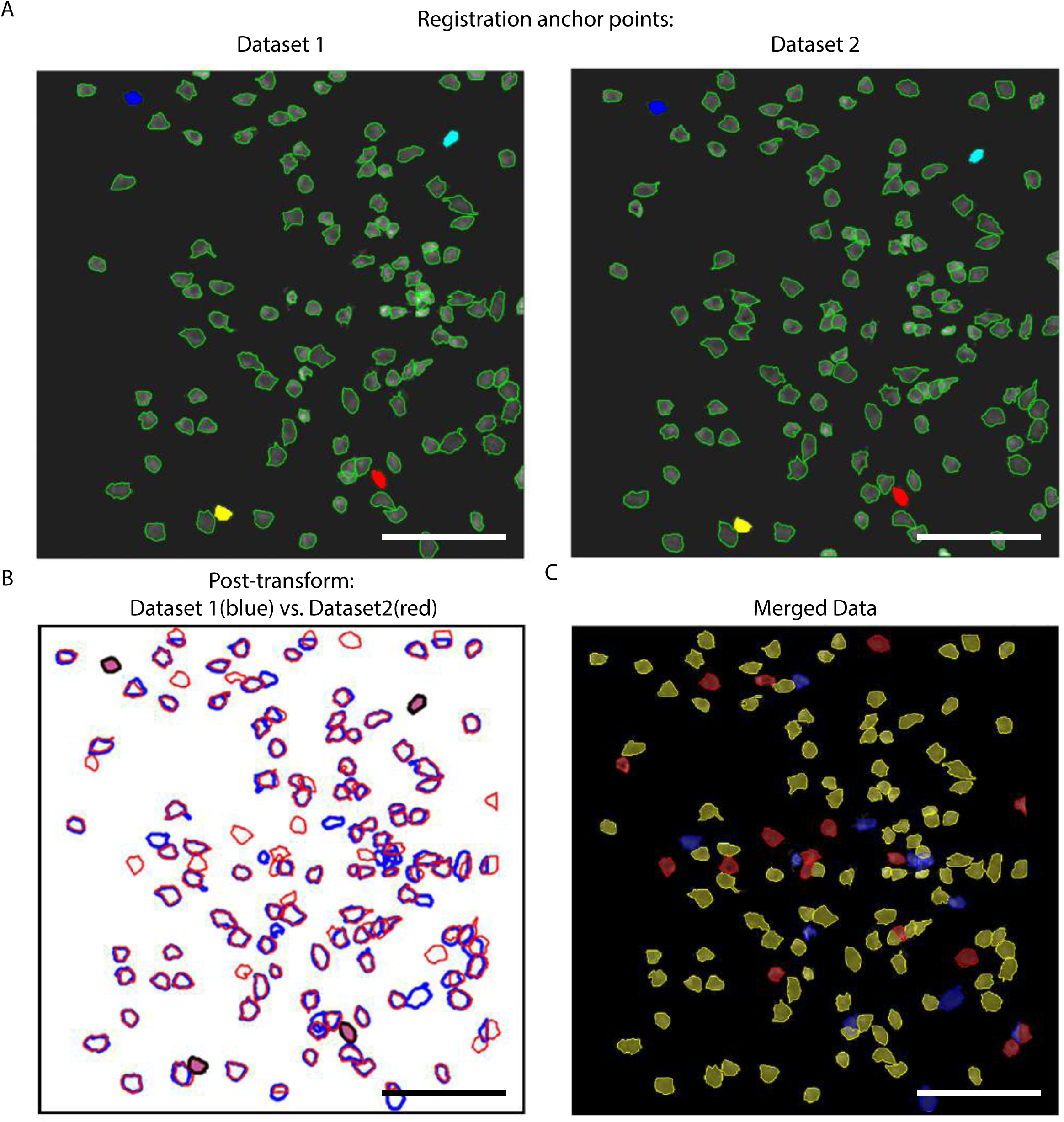
Cross-day neuronal alignment. **A,** Selection of 4 pairs of active neurons shown with different colors in both data sets (dark blue, light blue, yellow, and red filters). **B,** Post-transformation alignment of both fields of view (FOV). Blue filter contours denote active neurons from data set 1, and red filter contours denote active neurons from data set 2. Filled filters represent the original anchor points used in **A**.**C,** Merged FOV image. Blue filters indicate neurons active only in data set 1, red filters indicate neurons active only in data set 2, and yellow filters represent neurons that were active in both data sets. Scale bars: 100µm.

## REFERENCES

1. Albarran, E., Raissi, A., Jáidar, O., Shatz, C. J., & Ding, J. B. (2021). Enhancing motor learning by increasing the stability of newly formed dendritic spines in the motor cortex. Neuron, 109(20), 3298–3311.

2. Albarran, E., Sun, Y., Liu, Y., Raju, K., Dong, A., Li, Y., … & Ding, J. B. (2023). Postsynaptic synucleins mediate endocannabinoid signaling. Nature Neuroscience, 26(6), 997–1007.

3. Arber, S., & Costa, R. M. (2022). Networking brainstem and basal ganglia circuits for movement. Nature Reviews Neuroscience, 23(6), 342–360.

4. Athalye, V. R., Santos, F. J., Carmena, J. M., & Costa, R. M. (2018). Evidence for a neural law of effect. Science, 359(6379), 1024–1029.

5. Bailey, K. R., & Mair, R. G. (2006). The role of striatum in initiation and execution of learned action sequences in rats. Journal of Neuroscience, 26(3), 1016–1025.

6. Barnes, T. D., Kubota, Y., Hu, D., Jin, D. Z., & Graybiel, A. M. (2005). Activity of striatal neurons reflects dynamic encoding and recoding of procedural memories. Nature, 437(7062), 1158–1161.

7. Butz, M. V., Herbort, O., & Hoffmann, J. (2007). Exploiting redundancy for flexible behavior: unsupervised learning in a modular sensorimotor control architecture. Psychological Review, 114(4), 1015.

8. Cai, D. J., Aharoni, D., Shuman, T., Shobe, J., Biane, J., Song, W., … & Silva, A. J. (2016). A shared neural ensemble links distinct contextual memories encoded close in time. Nature, 534(7605), 115–118.

9. Calabresi, P., Picconi, B., Tozzi, A., & Di Filippo, M. (2007). Dopamine-mediated regulation of corticostriatal synaptic plasticity. Trends in neurosciences, 30(5), 211–219.

10. Calabresi, P., Picconi, B., Tozzi, A., Ghiglieri, V., & Di Filippo, M. (2014). Direct and indirect pathways of basal ganglia: a critical reappraisal. Nature neuroscience, 17(8), 1022–1030.

11. Cataldi, S., Lacefield, C., Shashaank, N., Kumar, G., Boumhaouad, S., & Sulzer, D. (2022). Decreased dorsomedial striatum direct pathway neuronal activity is required for learned motor coordination. eneuro, 9(5).

12. Cui, G., Jun, S. B., Jin, X., Pham, M. D., Vogel, S. S., Lovinger, D. M., & Costa, R. M. (2013). Concurrent activation of striatal direct and indirect pathways during action initiation. Nature, 494(7436), 238–242.

13. Denève, S., & Machens, C. K. (2016). Efficient codes and balanced networks. Nature neuroscience, 19(3), 375–382.

14. Dhawale, A. K., Smith, M. A., & Ölveczky, B. P. (2017). The role of variability in motor learning. Annual review of neuroscience, 40, 479–498.

15. Dhawale, A. K., Wolff, S. B., Ko, R., & Ölveczky, B. P. (2021). The basal ganglia control the detailed kinematics of learned motor skills. Nature neuroscience, 24(9), 1256–1269.

16. Freeze, B. S., Kravitz, A. V., Hammack, N., Berke, J. D., & Kreitzer, A. C. (2013). Control of basal ganglia output by direct and indirect pathway projection neurons. Journal of Neuroscience, 33(47), 18531–18539.

17. Gittis, A. H., & Kreitzer, A. C. (2012). Striatal microcircuitry and movement disorders. Trends in neurosciences, 35(9), 557–564.

18. Graybiel, A. M. (2005). The basal ganglia: learning new tricks and loving it. Current opinion in neurobiology, 15(6), 638–644.

19. Grillner, S. (2006). Biological pattern generation: the cellular and computational logic of networks in motion. Neuron, 52(5), 751–766.

20. Huber, D., Gutnisky, D. A., Peron, S., O’connor, D. H., Wiegert, J. S., Tian, L., … & Svoboda, K. (2012). Multiple dynamic representations in the motor cortex during sensorimotor learning. Nature, 484(7395), 473–478.

21. Hwang, E. J., Dahlen, J. E., Hu, Y. Y., Aguilar, K., Yu, B., Mukundan, M., … & Komiyama, T. (2019). Disengagement of motor cortex from movement control during long-term learning. Science advances, 5(10), eaay0001.

22. Hwang, F. J., Roth, R. H., Wu, Y. W., Sun, Y., Kwon, D. K., Liu, Y., & Ding, J. B. (2022). Motor learning selectively strengthens cortical and striatal synapses of motor engram neurons. Neuron, 110(17), 2790–2801.

23. Inan, H., Schmuckermair, C., Tasci, T., Ahanonu, B. O., Hernandez, O., Lecoq, J., … & Schnitzer, M. J. (2021). Fast and statistically robust cell extraction from large-scale neural calcium imaging datasets. BioRxiv, 2021-03.

24. Isomura, Y., Takekawa, T., Harukuni, R., Handa, T., Aizawa, H., Takada, M., & Fukai, T. (2013). Reward-modulated motor information in identified striatum neurons. Journal of Neuroscience, 33(25), 10209–10220.

25. Josselyn, S. A., & Tonegawa, S. (2020). Memory engrams: Recalling the past and imagining the future. Science, 367(6473), eaaw4325.

26. Jin, X., & Costa, R. M. (2010). Start/stop signals emerge in nigrostriatal circuits during sequence learning. Nature, 466(7305), 457–462.

27. Kawai, R., Markman, T., Poddar, R., Ko, R., Fantana, A. L., Dhawale, A. K., … & Ölveczky, B. P. (2015). Motor cortex is required for learning but not for executing a motor skill. Neuron, 86(3), 800–812.

28. Klaus, A., Martins, G. J., Paixao, V. B., Zhou, P., Paninski, L., & Costa, R. M. (2017). The spatiotemporal organization of the striatum encodes action space. Neuron, 95(5), 1171–1180.

29. Komiyama, T., Sato, T. R., O’Connor, D. H., Zhang, Y. X., Huber, D., Hooks, B. M., … & Svoboda, K. (2010). Learning-related fine-scale specificity imaged in motor cortex circuits of behaving mice. Nature, 464(7292), 1182–1186.

30. Kondapavulur, S., Lemke, S. M., Darevsky, D., Guo, L., Khanna, P., & Ganguly, K. (2022). Transition from predictable to variable motor cortex and striatal ensemble patterning during behavioral exploration. Nature communications, 13(1), 2450.

31. Koralek, A. C., Costa, R. M., & Carmena, J. M. (2013). Temporally precise cell-specific coherence develops in corticostriatal networks during learning. Neuron, 79(5), 865–872.

32. Krok, A. C., Maltese, M., Mistry, P., Miao, X., Li, Y., & Tritsch, N. X. (2023). Intrinsic dopamine and acetylcholine dynamics in the striatum of mice. Nature, 621(7979), 543–549.

33. Laubach, M., Wessberg, J., & Nicolelis, M. A. (2000). Cortical ensemble activity increasingly predicts behaviour outcomes during learning of a motor task. Nature, 405(6786), 567–571.

34. Lawrence, D. G., & Kuypers, H. G. (1968). The functional organization of the motor system in the monkey: I. The effects of bilateral pyramidal lesions. Brain, 91(1), 1–14.

35. Lee, J., & Sabatini, B. L. (2021). Striatal indirect pathway mediates exploration via collicular competition. Nature, 599(7886), 645–649.

36. Lemke, S. M., Ramanathan, D. S., Guo, L., Won, S. J., & Ganguly, K. (2019). Emergent modular neural control drives coordinated motor actions. Nature neuroscience, 22(7), 1122–1131.

37. Lemke, S. M., Ramanathan, D. S., Darevksy, D., Egert, D., Berke, J. D., & Ganguly, K. (2021). Coupling between motor cortex and striatum increases during sleep over long-term skill learning. Elife, 10, e64303.

38. Liljeholm, M., & O’Doherty, J. P. (2012). Contributions of the striatum to learning, motivation, and performance: an associative account. Trends in cognitive sciences, 16(9), 467–475.

39. Lindsey, J., Markowitz, J. E., Datta, S. R., & Litwin-Kumar, A. (2024). Dynamics of striatal action selection and reinforcement learning. bioRxiv, 2024-02.

40. Ma, L., Day-Cooney, J., Benavides, O. J., Muniak, M. A., Qin, M., Ding, J. B., … & Zhong, H. (2022). Locomotion activates PKA through dopamine and adenosine in striatal neurons. Nature, 611(7937), 762–768.

41. Maltese, M., March, J. R., Bashaw, A. G., & Tritsch, N. X. (2021). Dopamine differentially modulates the size of projection neuron ensembles in the intact and dopamine-depleted striatum. Elife, 10, e68041.

42. Markowitz, J. E., Gillis, W. F., Beron, C. C., Neufeld, S. Q., Robertson, K., Bhagat, N. D., … & Datta, S. R. (2018). The striatum organizes 3D behavior via moment-to-moment action selection. Cell, 174(1), 44–58.

43. Markowitz, J. E., & Datta, S. R. (2020). The striatum specifies the statistics of behavior. Neuropsychopharmacology, 45(1), 222.

44. Masamizu, Y., Tanaka, Y. R., Tanaka, Y. H., Hira, R., Ohkubo, F., Kitamura, K., … & Matsuzaki, M. (2014). Two distinct layer-specific dynamics of cortical ensembles during learning of a motor task. Nature neuroscience, 17(7), 987–994.

45. Mau, W., Hasselmo, M. E., & Cai, D. J. (2020). The brain in motion: How ensemble fluidity drives memory-updating and flexibility. Elife, 9, e63550.

46. McHaffie, J. G., Stanford, T. R., Stein, B. E., Coizet, V., & Redgrave, P. (2005). Subcortical loops through the basal ganglia. Trends in neurosciences, 28(8), 401–407.

47. Narayanan, N. S., Kimchi, E. Y., & Laubach, M. (2005). Redundancy and synergy of neuronal ensembles in motor cortex. Journal of Neuroscience, 25(17), 4207–4216.

48. Nigam, S., Pojoga, S., & Dragoi, V. (2019). Synergistic coding of visual information in columnar networks. Neuron, 104(2), 402–411.

49. Pérez-Ortega, J., Alejandre-García, T., & Yuste, R. (2021). Long-term stability of cortical ensembles. Elife, 10, e64449.

50. Peters, A. J., Chen, S. X., & Komiyama, T. (2014). Emergence of reproducible spatiotemporal activity during motor learning. Nature, 510(7504), 263–267.

51. Peters, A. J., Liu, H., & Komiyama, T. (2017). Learning in the rodent motor cortex. Annual review of neuroscience, 40, 77–97.

52. Pimentel-Farfan, A. K., Báez-Cordero, A. S., Peña-Rangel, T. M., & Rueda-Orozco, P. E. (2022). Cortico-striatal circuits for bilaterally coordinated movements. Science advances, 8(9), eabk2241.

53. Pisani, A., Centonze, D., Bernardi, G., & Calabresi, P. (2005). Striatal synaptic plasticity: implications for motor learning and Parkinson’s disease. Movement Disorders, 20(4), 395–402.

54. Pnevmatikakis, E. A., Soudry, D., Gao, Y., Machado, T. A., Merel, J., Pfau, D., … & Paninski, L. (2016). Simultaneous denoising, deconvolution, and demixing of calcium imaging data. Neuron, 89(2), 285–299.

55. Pnevmatikakis, E. A., & Giovannucci, A. (2017). NoRMCorre: An online algorithm for piecewise rigid motion correction of calcium imaging data. Journal of neuroscience methods, 291, 83–94.

56. Raman, D. V., Rotondo, A. P., & O’Leary, T. (2019). Fundamental bounds on learning performance in neural circuits. Proceedings of the National Academy of Sciences, 116(21), 10537–10546.

57. Rolls, E. T., & Treves, A. (2011). The neuronal encoding of information in the brain. Progress in neurobiology, 95(3), 448–490.

58. Roth, R. H., Cudmore, R. H., Tan, H. L., Hong, I., Zhang, Y., & Huganir, R. L. (2020). Cortical synaptic AMPA receptor plasticity during motor learning. Neuron, 105(5), 895–908.

59. Rueda-Orozco, P. E., & Robbe, D. (2015). The striatum multiplexes contextual and kinematic information to constrain motor habits execution. Nature neuroscience, 18(3), 453–460.

60. Santos, F. J., Oliveira, R. F., Jin, X., & Costa, R. M. (2015). Corticostriatal dynamics encode the refinement of specific behavioral variability during skill learning. Elife, 4, e09423.

61. Sheng, M. J., Lu, D., Shen, Z. M., & Poo, M. M. (2019). Emergence of stable striatal D1R and D2R neuronal ensembles with distinct firing sequence during motor learning. Proceedings of the National Academy of Sciences, 116(22), 11038–11047.

62. Tang, C., Pawlak, A. P., Prokopenko, V., & West, M. O. (2007). Changes in activity of the striatum during formation of a motor habit. European Journal of Neuroscience, 25(4), 1212–1227.

63. Tang, E., Mattar, M. G., Giusti, C., Lydon-Staley, D. M., Thompson-Schill, S. L., & Bassett, D. S. (2019). Effective learning is accompanied by high-dimensional and efficient representations of neural activity. Nature neuroscience, 22(6), 1000–1009.

64. Tecuapetla, F., Jin, X., Lima, S. Q., & Costa, R. M. (2016). Complementary contributions of striatal projection pathways to action initiation and execution. Cell, 166(3), 703–715.

65. Turner, R. S., & Desmurget, M. (2010). Basal ganglia contributions to motor control: a vigorous tutor. Current opinion in neurobiology, 20(6), 704–716.

66. Vicente, A. M., Galvão-Ferreira, P., Tecuapetla, F., & Costa, R. M. (2016). Direct and indirect dorsolateral striatum pathways reinforce different action strategies. Current Biology, 26(7), R267–R269.

67. Wilson, C. J. (2004). Basal ganglia. The synaptic organization of the brain, 5, 361–414.

68. Yang, R., Tuan, R. R. L., Hwang, F. J., Bloodgood, D. W., Kong, D., & Ding, J. B. (2023). Dichotomous regulation of striatal plasticity by dynorphin. Molecular psychiatry, 28(1), 434–447.

69. Yin, H. H. (2010). The sensorimotor striatum is necessary for serial order learning. Journal of Neuroscience, 30(44), 14719–14723.

70. Yin, H. H., Knowlton, B. J., & Balleine, B. W. (2004). Lesions of dorsolateral striatum preserve outcome expectancy but disrupt habit formation in instrumental learning. European journal of neuroscience, 19(1), 181–189.

71. Yin, H. H., Mulcare, S. P., Hilário, M. R., Clouse, E., Holloway, T., Davis, M. I., … & Costa, R. M. (2009). Dynamic reorganization of striatal circuits during the acquisition and consolidation of a skill. Nature neuroscience, 12(3), 333–341.

72. Yuste, R., Cossart, R., & Yaksi, E. (2024). Neuronal ensembles: Building blocks of neural circuits. Neuron.

